# Pumilio and Brain tumor regulate the expression of Smaug during the *Drosophila* maternal-to-zygotic transition

**DOI:** 10.64898/2026.08.19.745818

**Authors:** Alexander J. Marsolais, Mariana Kekis, Harrison W. Smith, Zhuyi Wang, Chalini Weerasooriya, Timothy R. Hughes, Howard D. Lipshitz, Craig A. Smibert

## Abstract

During the *Drosophila* maternal-to-zygotic transition, maternally deposited RNAs are cleared in a temporally controlled manner. The RNA-binding protein Smaug is a major regulator of maternal mRNA decay and its expression at both the RNA and protein level is limited to a narrow temporal window during the MZT. Here we show that Pumilio RNA-binding protein promotes degradation of *smaug* mRNA at the end of the MZT through Pumilio-binding cis-elements in the *smaug* 3’ untranslated region, and that disruption of this regulation leads to ectopic Smaug protein expression beyond its normal developmental window. We further find that another RNA-binding protein, Brain tumor, also contributes to repression of ectopic Smaug expression. Transcriptome-wide analyses of embryos lacking Pumilio reveal that it directly regulates hundreds of maternal mRNAs after zygotic genome activation, with many of these Pumilio targets also regulated by Brain tumor. The ectopic Smaug protein that results from loss of either Pumilio or Brain tumor causes widespread downregulation of mRNAs that contain Smaug binding sites. Together, these findings define a post-transcriptional regulatory pathway in which Pumilio and Brain tumor ensure orderly progression of the *Drosophila* maternal-to-zygotic transition by clearing *smaug* mRNA and preventing ectopic Smaug activity.

## INTRODUCTION

Early animal embryos are transcriptionally silent and, thus, the early phases of development are directed by maternally deposited gene products that are synthesized during oogenesis and deposited into the developing egg (Vastenhouw *et al*. 2019; Kojima *et al*. 2025). As embryogenesis proceeds, transcription initiates in the embryo while many maternal mRNAs and a smaller subset of maternal proteins are degraded; thus, control of development is transferred from maternal products to those encoded by the zygote. This process is known as the maternal-to-zygotic transition (MZT).

Much of our understanding of the MZT comes from work in *Drosophila* (Harrison *et al*. 2023). For example, the existence of temporally distinct pathways of mRNA decay was first recognized in the fly (Bashirullah *et al*. 1999). Some of these pathways initiate early in embryogenesis and require only maternally contributed products, while others initiate later and require one or more zygotically produced factors. Hereafter, we will refer to these temporally distinct decay programs as ‘early’ and ‘late’ pathways. A major driver of early decay in the fly embryo is the RNA-binding protein, Smaug (SMG). SMG binds to target mRNAs through stem-loop structures known as SMG recognition elements (SREs) (Smibert *et al*. 1996; Aviv *et al*. 2003; Semotok *et al*. 2005; Aviv *et al*. 2006; Tadros *et al*. 2007; Semotok *et al*. 2008; Laver *et al*. 2015b) and recruits factors that repress target mRNA expression, including the CCR4/NOT deadenylase complex (Nelson *et al*. 2004; Semotok *et al*. 2005; Semotok *et al*. 2008; Jeske *et al*. 2011; Pinder and Smibert 2013; Gotze *et al*. 2017). CCR4/NOT recruitment results in poly(A) tail shortening which triggers RNA degradation.

SMG protein is expressed in a narrow temporal window (Smibert *et al*. 1996; Smibert *et al*. 1999; Tadros *et al*. 2007; Benoit *et al*. 2009). During oogenesis, SMG protein is undetectable as its cognate mRNA is translationally repressed. Upon egg activation, this repression is relieved, and the protein accumulates in the early embryo. Towards the end of the MZT, SMG is degraded by a Skp1-Cullin-F-box (SCF) complex (Cao *et al*. 2020). The timing of this degradation involves the F-box protein, Bard, which is synthesized from a zygotically transcribed mRNA (Cao *et al*. 2020; Cao *et al*. 2022). Bard itself is tightly regulated with both the mRNA and protein rapidly cleared at the end of the MZT.

Abrogation of SMG degradation by preventing its interaction with SCF, results in persistent SMG expression and consequent downregulation of mRNAs that carry SREs (Cao *et al*. 2020). These results highlight that a properly orchestrated MZT likely requires mechanisms to turn off specific pathways of mRNA decay to ensure that zygotically synthesized RNAs are not inappropriately targeted for degradation.

SMG protein and mRNA are cleared from the embryo over a similar time frame (Smibert *et al*. 1996; Smibert *et al*. 1999; Tadros *et al*. 2007; Benoit *et al*. 2009). In the earlier studies of SMG protein clearance (Cao *et al*. 2020; Cao *et al*. 2022), when SCF function was abrogated and SMG protein persisted, *smg* mRNA degradation proceeded as in wild type. Those experiments, therefore, were unable to assess the role of *smg* transcript degradation *per se* during the MZT. Here, we set out to identify the *cis* and *trans*-acting factors that function in *smg* mRNA decay. We show that the RNA-binding protein Pumilio (PUM) triggers *smg* mRNA decay via PUM-binding elements (PBEs) in the *smg* 3’ UTR. Disrupting PUM-mediated decay results in the expression of SMG protein beyond its normal temporal window. Our previous work indicated that *smg* mRNA is targeted by the RNA-binding protein Brain tumor (BRAT) (Laver *et al*. 2015b) and here we show that BRAT is required to repress ectopic SMG protein expression. We use loss-of-function *pum* alleles to demonstrate that PUM has a direct role in degrading hundreds of mRNAs after the onset of zygotic transcription. Many of these transcripts are degraded in a BRAT-dependent manner late in the MZT, consistent with extensive co-regulation by these two RNA-binding proteins. Strikingly, the ectopic SMG protein that is expressed in the absence of PUM or BRAT has a major effect on the embryo’s transcriptome by degrading mRNAs that carry SREs. Thus, downregulation of *smg* mRNA by PUM and BRAT is required for an orderly progression of the transcriptome through the MZT. The fact that both SMG protein and *smg* mRNA must be cleared to permit an orderly MZT emphasizes the key role of both post-transcriptional and post-translational mechanisms in this developmental process.

## MATERIALS AND METHODS

### *Drosophila* stocks

The following stocks were used: *pum^7^*/TM3, Sb (Forbes and Lehmann 1998), *pum^Msc^*/TM3, Sb (Barker *et al*. 1992), *brat^fs1^/*CyO (Schupbach and Wieschaus 1991) and *Df(2L)TE37C-7/*CyO (Stathakis *et al*. 1995). Embryos lacking wild-type maternal PUM were collected from *pum^7^/pum^Msc^*mothers, while embryos lacking wild-type maternal BRAT were collected from *brat^fs1^/Df(2L)TE37C-7* mothers. Wild-type flies were *w^1118^*. Flies were cultivated at 25 °C under standard laboratory conditions.

Tagging of the endogenous *pum* gene at the 3’ end of its open-reading frame (ORF) with a 3xFLAG sequence, which tags all known *pum* isoforms, was performed by Well Genetics [13F.-12, No. 93, Sec. 1, Xintai 5th Rd., Xizhi Dist., New Taipei City 221416, Taiwan (R.O.C.)].

### Transgenic reporters

Reporters were constructed in modified pCaSpeR vectors containing an attB site for site-directed transgenesis using the attP-attB-φC31 integrase (Bischof *et al*. 2007). All constructs were inserted at the attP40 site on the *Drosophila* second chromosome.

Note that the SGS transgene described here is not identical to the SGS construct we previously described (Tadros *et al*. 2007). Constructs were either driven by the *αTub84B* or *smg* promoters cloned as AscI to NotI fragments. All *αTub84B* promoter-driven reporters retained the endogenous *αTub84B* intron which has been shown to contain promoter/enhancer elements (O’Donnell *et al*. 1994), and which lies immediately downstream of the *αTub84B* 5’ UTR and ATG start codon. The fragment containing the *αTub84B* promoter, 5’ UTR, start codon and intron was amplified from genomic DNA isolated as a ∼ 2.7 kb fragment using primers (forward primer – 5’-CTTACCGATGTCGACGAAGAGG-3’ and reverse primer 5’-CTGTGGATGAGGAGGAAGGGA-3’). The endogenous *αTub84B* start codon was eliminated through QuikChange PCR using primers (sense primer – 5’-TTCCAATAAAAACTCAATGTGGTGAGTACTTTAAAAAAA-3’ and anti-sense primer – 5’-TTTTTTTAAAGTACTCACCACATTGAGTTTTTATTGGAA-3’). All *smg* promoter-driven constructs retained the endogenous intron located in the *smg* isoform RA 5’ UTR. The *smg* promoter and 5’ UTR fragment is approximately 6.5 kb and was amplified from genomic DNA.

Reporters contained either the *eGFP* open reading frame cloned as NotI/SwaI fragment downstream of the *αTub84B* or *smg* promoter fragments. Downstream of the open reading frame, either the *αTub84B* or *smg* 3’ UTRs and downstream genomic sequence were cloned as SwaI/SbfI fragments. The *αTub84B* 3’ UTR fragment is 815 nucleotides long and includes the 287 nucleotide *αTub84B* 3’ UTR and 561 nucleotides of downstream genomic sequence. The *smg* fragment is 2217 nucleotides long and includes the *smg* 3’ UTR and approximately 1 kilobase of downstream genomic sequence.

To clone fragments of the *smg* 3’ UTR into *TGT* reporters, the *αTub84B* 3’ UTR was further modified to include a XhoI site upstream of the BamHI site, and fragments of the *smg* 3’ UTR were amplified with XhoI and BamHI sites and cloned. PBE-versions of *smg* 3’ UTR-bearing reporters were generated through overlap extension PCR to mutate each TGTA in the *smg* 3’ UTR to ACAA.

### Embryo collections and processing

Embryos were collected for 1, 1.5 or 2 hours on yeasted apple juice plates, aged appropriately, and dechorionated with 4% hypochlorite for 2 minutes, followed by extensive washing with 0.1% Triton X-100. Total RNA was then purified using TRI-reagent (Bioshop) according to the manufacturer’s instructions.

For immunoprecipitation (IP) experiments 0-2 and 2-4 hour (h) embryos were disrupted in IP lysis buffer (18 mL/gram of 1M trehalose, 150mM KCl, 30mM HEPES-KOH pH 7.4, 1mM MgCl_2_, 1mM AEBSF, 2 mM benzamidine, 2 μg/mL leupeptin, 2 μg/mL pepstatin, 10 mM beta-glycerolphosphate, 1 mM sodium pyrophosphate, 5 mM NaF, 1 mM sodium orthovanadate and 1 mM DTT). Extracts were clarified by centrifugation at 21,000g for 15 minutes, and the resulting supernatants were stored at −80°C.

For Western blots embryos were lysed in Western lysis buffer (150 mM KCl, 30 mM HEPES-KOH pH 7.4, 1mM AEBSF, 2 mM benzamidine, 2 μg/mL leupeptin, 2 μg/mL pepstatin). Extracts were clarified by centrifugation at 21,000g for 15 minutes, and the resulting supernatants was stored at −80°C.

### RNA analysis

For Northern blots, 4 µg of total RNA was resolved on 1% agarose denaturing gels as described in (Selden 1987), transferred to nitrocellulose and incubated with ^32^P-labelled probes generated via random priming using the Prime-a-Gene Labeling System (Promega). Membranes were imaged using Phosphorimager Screens (GE), which were scanned using a Typhoon scanner (GE). Band intensities were quantified using ImageJ software (Schneider *et al*. 2012).

For IPs, frozen extracts were thawed, diluted IP lysis buffer to a protein concentration of 3mg/mL and Triton X-100 was added to a final concentration of 0.1%. The extract was then centrifuged at 21,000g for 15 minutes. Approximately 1.3 mL of supernatant was then mixed end-over-end with 30µl of Anti-FLAG M2 Affinity Gel (Millipore Sigma) for 3 hours and, after extensive washing in IP lysis buffer, RNA was purified from the beads as described above. RT-qPCR was performed as described in (Low *et al*. 2026) using random hexamers to prime cDNA synthesis.

For microarray analysis cDNA synthesis was performed with random primers as described (Kapranov *et al*. 2007). Custom Agilent microarrays (GEO accession GPL17421) were used containing ∼44,000 probes, representing 12,396 genes from *Drosophila melanogaster* release 3.41. The array was designed using OligoPicker software (Wang and Seed 2003). Microarray slides were scanned with an Agilent High-Resolution C Scanner and quantified using ImaGene software.

Raw data was then normalized using robust multi-array average (RMA) (Bolstad *et al*. 2003; Irizarry *et al*. 2003) and log2-transformed using the ArrayStar program (DNASTAR, Inc. Madison, WI). Normalized data were then subjected to one-class significance of microarray (SAM) analysis (Tusher *et al*. 2001) using the Multi-experiment Viewer (MeV) software (Saeed *et al*. 2003; Saeed *et al*. 2006). Any gene having an FDR < 5% in this analysis in at least one time point for at least one genotype was defined as ‘expressed’. The normalized data for each of these genes was compared in wild-type and *pum* mutant embryos using a two-class SAM analysis in the MeV software package. Differentially expressed genes were defined as those with an FDR < 5% and fold difference in wild type versus *pum* mutant of > 1.5 or < −1.5-fold. FlyBase gene numbers for each gene in the dataset were then updated using the gProfiler2 v0.2.3 R package (Kolberg *et al*. 2020) to Flybase version 6.58 (Ozturk-Colak *et al*. 2024). Where necessary the same method was used to convert FlyBase gene numbers from previously published datasets to Flybase version 6.58 to facilitate comparisons with our *pum* mutant transcriptome analysis.

### Western blots

Western blots employed the following primary antibodies: mouse anti-β-tubulin E7 and mouse anti-chicken actin JLA20 (both from the Developmental Studies Hybridoma Bank, Iowa City), and guinea pig anti-SMG (Tadros *et al*. 2007). HRP-conjugated secondary antibodies were purchased from Jackson ImmunoResearch. Westerns were visualized using an Immobilon Luminata Crescendo Western HRP substrate (Millipore), imaged using a BioRad ChemiDoc MP Universal Hood III imager, and band intensities were quantified using ImageJ software (Schneider *et al*. 2012).

### Motif analysis

For PBE score determination, we searched the longest transcript isoform for each gene for matches to the sequence UGUANA, the PBE determined by RNAcompete (Ray *et al*. 2013) and assessed each match’s accessibility using on RNAplfold (ViennaRNA-2.5.1) (Hofacker *et al*. 1994; Lorenz *et al*. 2011). The RNAplfold results were then analyzed using the MFRA (Motif and Flanking Region Accessibility) pipeline (Hu *et al*. 2026). Briefly, each transcript was scanned with RNAplfold using an 80-nucleotide window, and the accessibility of each motif was defined as the probability that it is single-stranded and thus available for protein binding. Finally, a PBE score for each transcript was calculated by summing the accessibility of all predicted matches. Transcript SRE scores for the longest transcript isoform for each gene are from Siddiqui *et al*. (2024).

## RESULTS

### The *smg* 3’ UTR is the major driver of transcript decay

To explore the mechanisms that regulate *smg* mRNA decay, we first mapped the *cis*-acting elements within the *smg* transcript that function in its degradation. To do so we used a transgenic reporter construct that carries the α*Tub84B* promoter driving the expression of an mRNA with the α*Tub84B* 5’ UTR, the *GFP* ORF, and the α*Tub84B* 3’ UTR, which we refer to as *TGT*, where the letters indicate, respectively, the source of the promoter+5’ UTR, ORF, and 3’ UTR (Fig. 1a).

**Figure 1.**
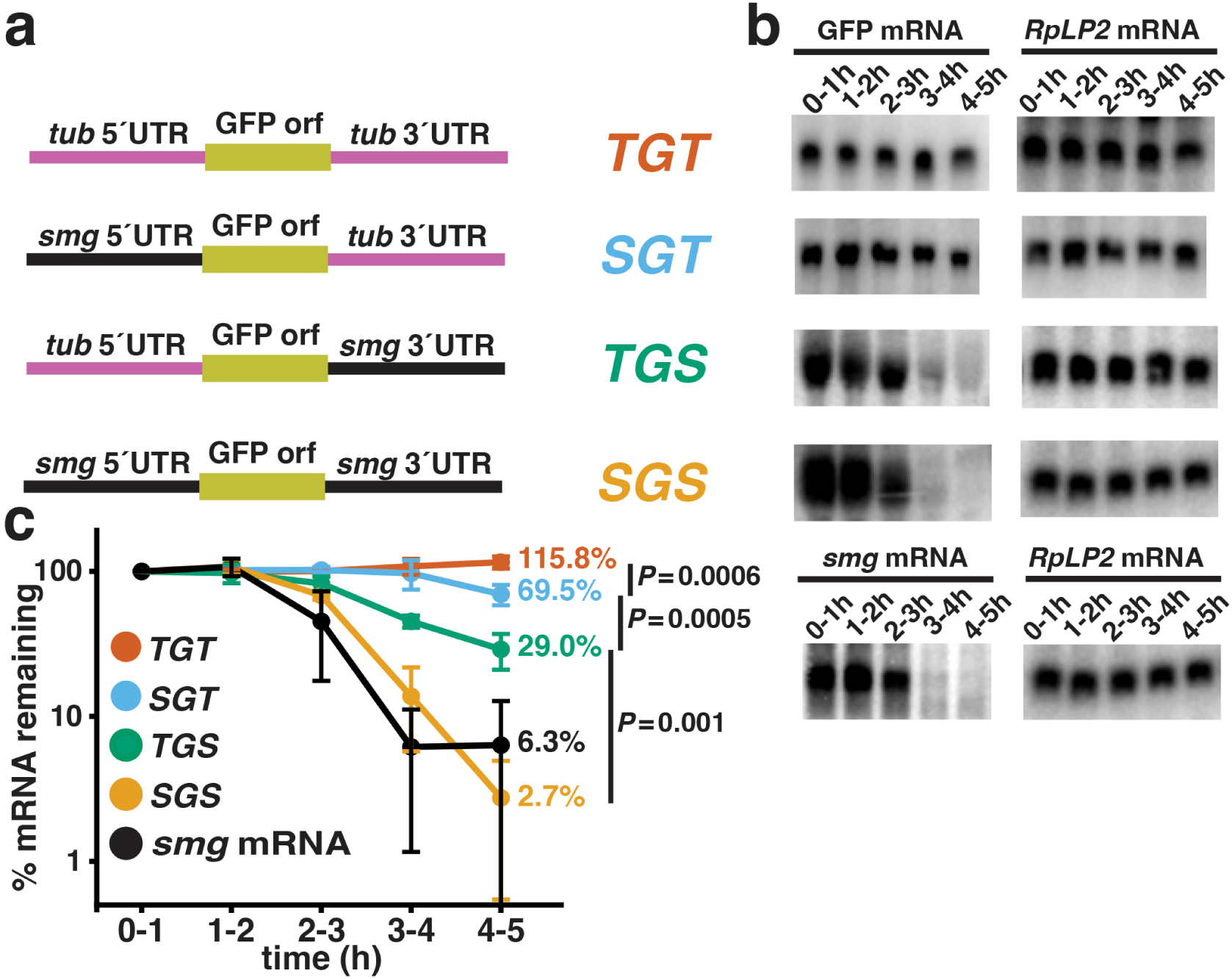
*smg* mRNA decay is driven by the *smg* 3’ UTR. (a) The transgenic reporters generated to assess the role of the *smg* 5’and 3’ UTRs in *smg* mRNA degradation are diagrammed. Each construct is given a three-letter designation where the first letter and last letter indicate the origin of the reporter’s promoter-5’ UTR or the 3’ UTR-downstream genomic sequences, respectively (i.e. S for *smg* and T for α*Tub84B*), while the middle letter for all is a G, indicating that each carries the GFP ORF. (b) Total RNA was harvested from embryos expressing the transgenic mRNAs described in (a) at the indicated time intervals and subjected to Northern blot analysis using a GFP probe. *RpLP2* mRNA served as a loading control. The degradation of endogenous *smg* mRNA was assessed using RNA samples collected from embryos expressing the *TGT* transgene using a *smg*-specific probe. (c) After normalizing transgenic mRNA levels using the loading control, the amount in 0-1 h embryos was set to 100%. N=3 for SGS and 4 for all others. Error bars represent standard deviation. The percent remaining of each mRNA at the last time point is indicated, while one-tailed Student’s t-test *P* values compare the levels of transgenic mRNA remaining at the last time point in *SGS* versus *TGS*, *TGS* versus *SGT* and *SGT* versus *TGT*.

*TGT* mRNA was stable over the first five hours of embryogenesis, as assayed by Northern blot (Fig. 1b and c). Replacing the *tubulin* promoter, 5’ and 3’ UTR with the *smg* promoter, 5’ and 3’ UTR, to give SGS, resulted in an mRNA that was degraded with a profile similar to endogenous *smg* mRNA. Switching the 3’ UTR sequences to give *TGS* resulted in an mRNA that was degraded, albeit not to the same extent as endogenous *smg* or *SGS*. *SGT* mRNA, where the *tubulin* promoter and 5’ UTR were replaced with the *smg* promoter and 5’ UTR, was stable through the stages during which endogenous *smg*, SGS and *TGS* transcripts underwent degradation (2-3 and 3-4 h), then was partially degraded during the fifth hour of embryogenesis. At the 4-5 h time point there was significantly more *TGS* mRNA relative to *SGS* mRNA, *SGT* mRNA relative to *TGS* mRNA and *TGT* mRNA relative to *SGT* mRNA (one-tailed Student’s *t*-test *P* values 0.001, 0.0005 and 0.0006, respectively). Taken together, these data suggest that the *smg* 3’ UTR makes a major contribution to the degradation of *smg* RNA, while the *smg* 5’ UTR plays a smaller role.

The timing of *smg* mRNA degradation is consistent with a requirement for zygotic transcription, which initiates at high levels at ∼2.5 hours. Furthermore, northern blot analysis of *smg* mRNA in unfertilized eggs confirmed that it is stable in the absence of zygotic transcription (Supplemental Fig. 1). These data are consistent with previous transcriptomic analyses, which indicated that *smg* mRNA decay requires zygotic transcription (Tadros *et al*. 2007; Thomsen *et al*. 2010).

### PUM binds to and directs *smg* mRNA decay

A previous study found that *smg* mRNA copurifies with a tagged transgenic version of the PUM RNA-binding domain isolated from either ovaries or 0-16 h embryos (Gerber *et al*. 2006). To assess whether *smg* transcripts copurify with endogenous, full-length PUM, we employed a fly line where the *pum* gene carried three FLAG epitopes inserted at the 3’ end of the *pum* open reading frame. FLAG IPs were carried out from 0-2 and 2-4 h embryo extracts and the levels of *smg* mRNA that co-purified with PUM were quantified via RT-qPCR. *smg* mRNA was highly enriched in FLAG IPs compared to similar purifications from extracts lacking FLAG-tagged PUM (Fig. 2a), while an irrelevant mRNA, *RpL32*, was not.

**Figure 2.**
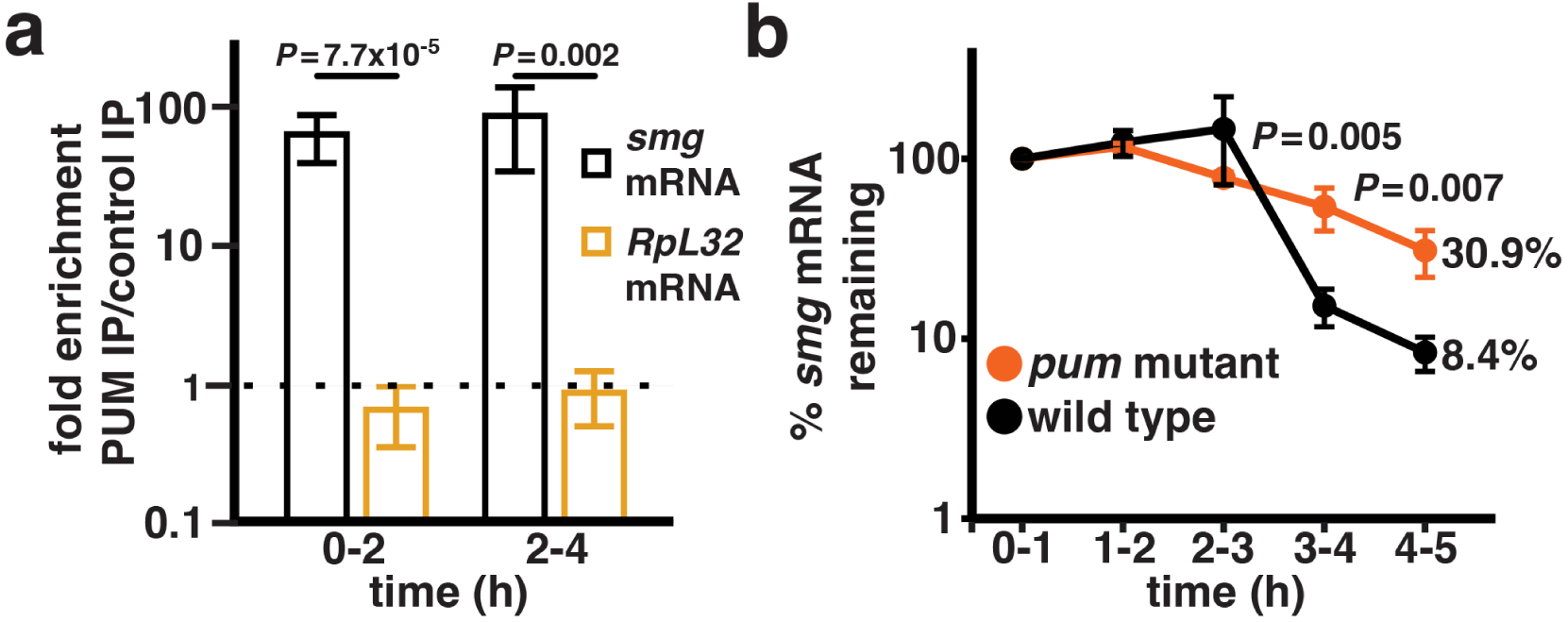
*smg* mRNA copurifies with PUM protein and is stabilized in a *pum* mutant. (a) 0-2 and 2-4 h embryo extracts, where endogenous PUM is tagged at its C-terminus with a 3xFLAG, were subjected to IP with anti-FLAG beads. The levels of *smg* mRNA and *RpL32* mRNA present in the IPs were then determined using RT-qPCR. Fold enrichment was calculated by dividing the amount of each mRNA recovered from FLAG IPs containing tagged PUM by the amount recovered from control FLAG IPs performed using extracts lacking FLAG-tagged PUM. The dashed black line indicates one-fold enrichment (i.e., no enrichment). N=6, error bars indicate standard deviation, while two-tailed Student’s t-test *P* values assess the significance of the fold enrichment of *smg* mRNA relative to that of *RpL32* mRNA. (b) Total RNA was harvested from wild-type and *pum* mutant embryos at the indicated time intervals, and levels of *smg* mRNA were determined using RT-qPCR. After normalizing *smg* mRNA levels to the control *RpL32* levels, the amount in 0-1 h embryos was set to 100%; the percent remaining of each mRNA at the last time point is indicated. N=3. One-tailed Student’s t-test *P* values compare the levels of *smg* mRNA in wild-type versus *pum* mutant embryos at the last two time points.

To assess whether PUM is necessary for clearance of *smg* mRNA we used females with a heteroallelic combination of two strong *pum* mutant alleles: *pum^7^* (also known as *pum^ET7^*, a nonsense allele that truncates PUM prior to its RNA-binding domain) (Forbes and Lehmann 1998) and *pum^Msc^*(a chromosomal inversion that also removes the PUM RNA-binding domain) (Barker *et al*. 1992). Total RNA was harvested from 0-1, 1-2, 2-3, 3-4 and 4-5 h embryos collected from *pum^7^*/*pum^Msc^* mutant mothers (hereafter referred to as *pum* mutant embryos) and subjected to RT-qPCR analysis. Figure 2b shows that *smg* mRNA is significantly stabilized in *pum* mutant embryos compared to wild type.

We next searched the *smg* UTRs for potential PUM-binding elements (PBEs). PUM binds to 8mer sequences that begin with an invariant UGUA followed by four more variable nucleotides (Wang *et al*. 2002; Gerber *et al*. 2006; Ray *et al*. 2013; Laver *et al*. 2015a; Weidmann *et al*. 2016). To assess the relative affinity of potential PBEs within *smg* mRNA UTRs, we used RNAcompete data (Ray *et al*. 2013). RNAcompete involves incubating a protein of interest with an excess of a complex pool containing approximately 240,000 30-41mer RNAs. After purification of the protein in question, co-purifying RNAs are identified using microarrays. These assays are done in excess RNA, and therefore the amount of any RNA that is bound to the query protein is a relative measure of the RNA’s affinity for that protein. We used RNAcompete data that ranked each of the possible 256 UGUANNNN 8mers (Ray *et al*. 2013), based on their predicted affinity for PUM, with rank 1 representing the highest affinity site. The *smg* 5’ UTR carries a single UGUA sequence, while the 3’ UTR carries 12. The 5’ UTR site has an RNAcompete rank of 2, while the motifs in the *smg* 3’ UTR have ranks between 26 and 239 (Fig. 3a and Supplemental Table 1). All 12 of the 3’ UTR sites are located downstream of nucleotide 466 of the *smg* 3’ UTR, which has a total length of 1179 nucleotides. Given the dominant role of the 3’ UTR in *smg* mRNA degradation we chose to focus on the PBEs located there. To do this we inserted nucleotides 1-907 of the *smg* 3’ UTR into the *TGT* reporter construct to generate *TGT+smg3’UTR^PBE+^*. Nucleotide 907 is just upstream of a potential AAUAAA polyadenylation signal within the *smg* mRNA, and truncation at 907 ensured that the reporter mRNA would terminate at the poly(A) site within tubulin 3’ UTR to avoid generating transcripts that had different 3’ ends. The 907 nt fragment contains nine of the 12 aforementioned PBEs, lacking three with RNAcompete ranks of 239, 173 and 75.

**Figure 3.**
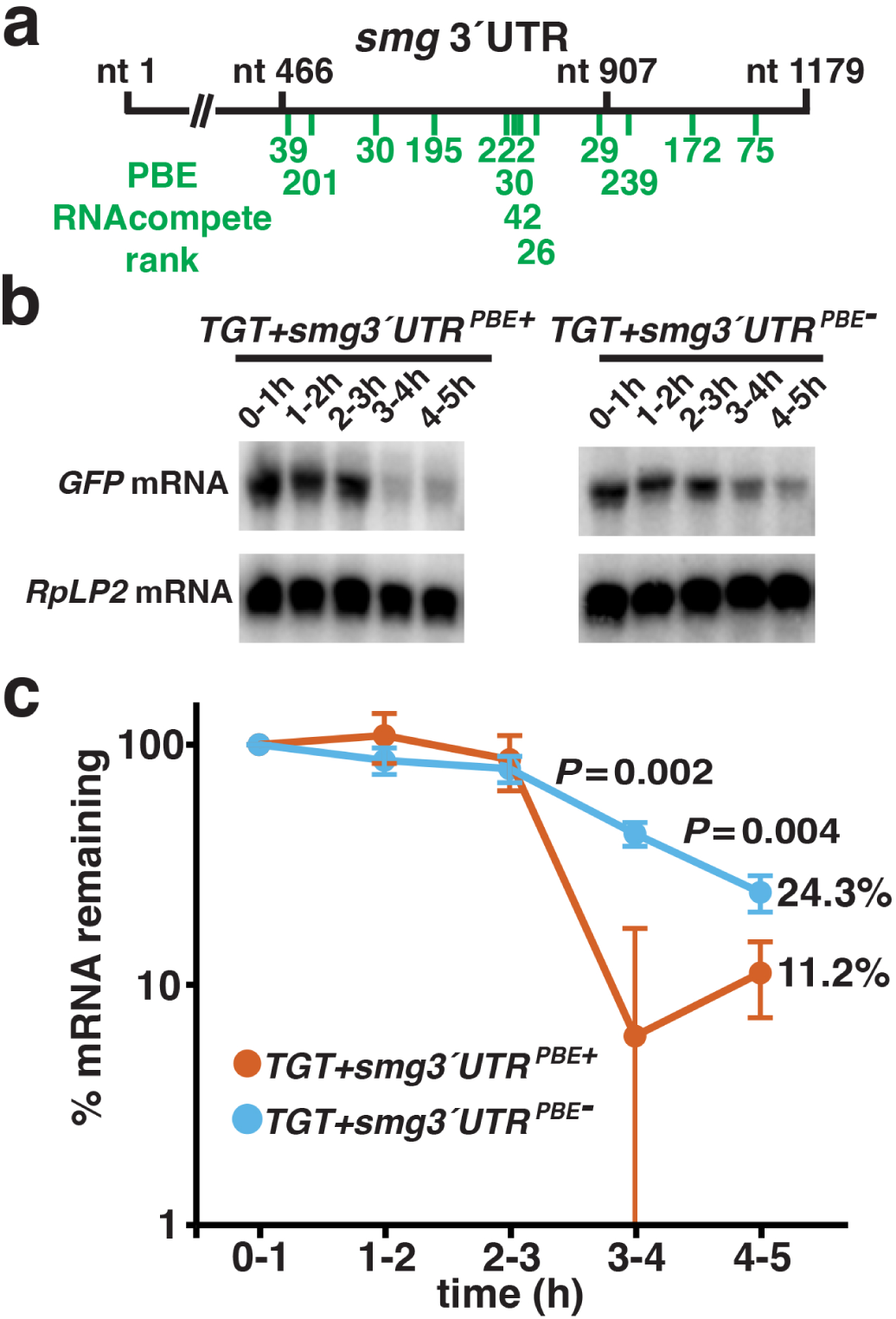
The PBEs in the *smg* 3’ UTR are required to destabilize a reporter mRNA. (a) The *smg* 3’ UTR is diagrammed, indicating the location of potential PBEs, using the sequence UGUANNNN. The rank of each site in the RNAcompete data is indicated, with rank 1 corresponding to the site with the highest apparent affinity for PUM. (b) Two transgenic *TGT* reporters were generated, where nucleotides 1-907 of the *smg* 3’ UTR were inserted into the α*Tub84B* 3’ UTR sequence of the reporter. *TGT+smg3*’UTR*^PBE+^* carries the wild-type *smg* 3’ UTR, which contains nine potential PBEs, while each of these sites is mutated from TGTA to ACAA in the *TGT+smg3*’*UTR^PBE-^*construct. Total RNA was harvested from transgenic embryos, and the levels of reporter mRNA were assessed using a GFP probe, where *RpLP2* mRNA served as a loading control. (c) After normalizing reporter mRNA levels to the loading control, the amount in 0-1 h embryos was set to 100%. N=4 for the *TGT+smg3’UTR^PBE+^*construct and N=3 for the *TGT+smg3’UTR^PBE-^* construct. Error bars represent standard deviation, and the percent remaining of each mRNA at the last time point is indicated. One-tailed Student’s *t*-test *P* values assess the significance of the difference between the levels of *TGT+smg3*’*UTR^PBE+^*and *TGT+smg3*’*UTR^PBE-^* mRNAs at the last two time points.

Northern blot analysis showed that insertion of the 907-nucleotide fragment destabilized the reporter mRNA to a similar extent and with a similar profile to endogenous *smg* mRNA, while mutation of all nine potential PBEs in construct *TGT+smg3’UTR^PBE-^* resulted in significant stabilization relative to the *TGT+smg3’UTR^PBE+^* RNA (Fig. 3b and c). Taken together, these data are consistent with a direct role for PUM in the degradation of *smg* mRNA.

### SMG protein persists in *pum* mutants

Having shown that PUM binds to and destabilizes the *smg* mRNA, we next asked whether the increase in *smg* transcript levels in *pum* mutants led to increased SMG protein. Western blots of SMG protein in wild-type and *pum* mutant embryos confirmed that SMG protein was cleared from wild-type embryos at 3-4 hours (Fig. 4) as previously reported (Benoit *et al*. 2009; Cao *et al*. 2020). In contrast, in *pum* mutant embryos, SMG protein levels were significantly upregulated in 3-4 and 4-5 h embryos. Together these data suggest that PUM-induced degradation of *smg* mRNA prevents ectopic expression of SMG protein late in the MZT.

**Figure 4.**
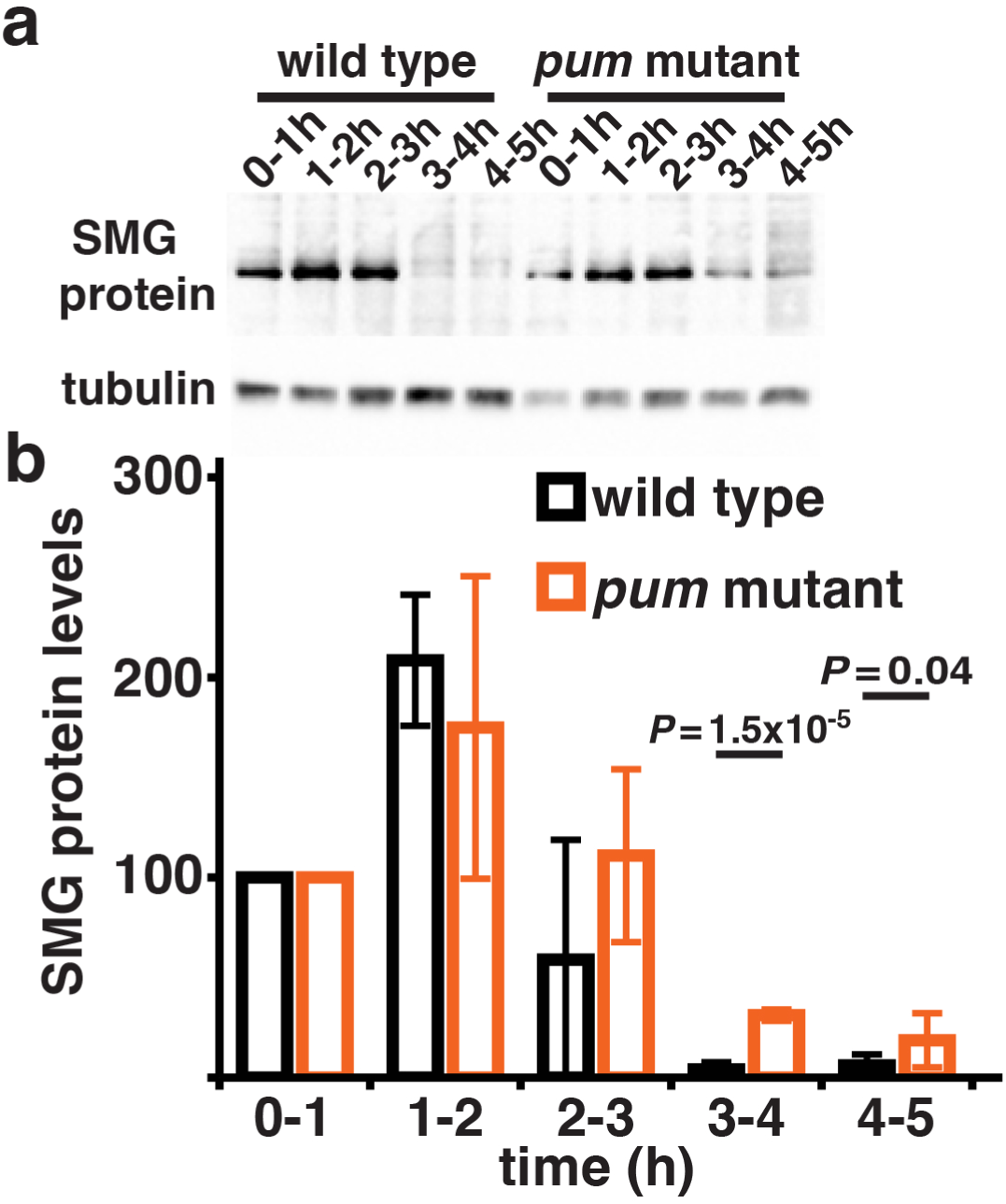
Ectopic SMG protein is detected in *pum* mutant embryos. (a) Total protein was harvested from wild-type and *pum* mutant embryos at the indicated time intervals, and levels of SMG protein were assayed via western blot. Blotting for tubulin served as the loading control. (b) After normalizing SMG protein levels to the loading control, the amount in 0-1 h embryos was set to 100%. N=4, with error bars representing standard deviation. One-tailed Student’s *t*-test *P* values assess the significance of the difference in the levels of SMG protein in wild-type versus *pum* mutant embryos at the last two time points.

### PUM is a major regulator of mRNA clearance during the late MZT

Previous work found that PUM-bound RNAs are enriched in transcripts that are unstable during the MZT (De Renzis *et al*. 2007; Thomsen *et al*. 2010; Laver *et al*. 2015a), suggesting that PUM directs a large-scale mRNA degradation pathway during this developmental phase. However, a comparison of the transcriptomes of wild-type females to *pum* mutant females that carried a weak *pum* mutant allele identified only a modest number of upregulated mRNAs (Gerber *et al*. 2006). To assess the effect of loss of *pum* function on the embryonic transcriptome, we harvested RNA from 0-1, 1-2, 2-3, 3-4 and 4-5 h embryos collected from *pum^7^*/*pum^Msc^* mutant mothers and used microarrays to compare their transcriptome to that of wild-type embryos. SAM (Tusher *et al*. 2001) was used to identify mRNAs that were significantly up-regulated in *pum* mutant embryos compared to wild-type at each time-point (Figure 5a; orange and blue dots indicate significantly upregulated or downregulated transcripts, respectively, with a > 1.5-fold change and a < 5% false discovery rate). Transcripts from only 20 genes were upregulated in 0-1, 1-2, and/or 2-3 h embryos, consistent with only a very minor role for PUM in the early to mid-MZT (Fig. 5a and b, Supplemental File 1). In contrast, in 3-4 and 4-5 h embryos we found 233 and 379 upregulated genes, respectively, suggesting a major role in transcript clearance late in the MZT. The list of genes up-regulated in 3-4 and 4-5 h embryos overlapped significantly (Fisher’s exact test *P*=1.41×10^-99^, odds ratio=26.81, Supplemental Table 2). The union of the 3-4 and 4-5 timepoints produced a list of 479 upregulated genes. Upregulated mRNAs included *hunchback* (*hb*), a well-characterized direct PUM target (Murata and Wharton 1995) and *bicoid*, an mRNA previously shown to be stabilized in *pum* mutant embryos (Gamberi *et al*. 2002) as well as *smg* mRNA, consistent with the results in Figures 2 and 3.

**Figure 5.**
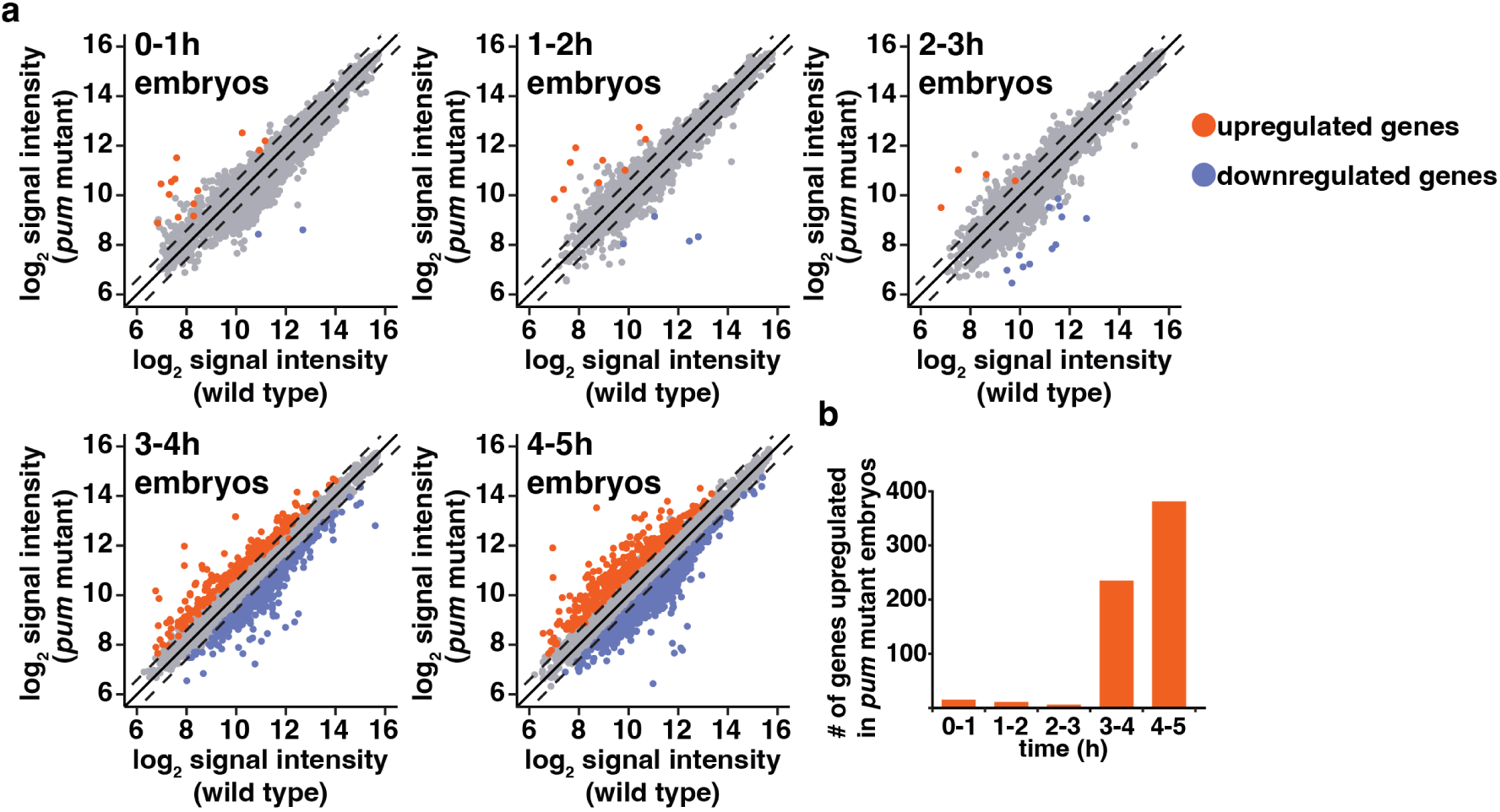
PUM regulates hundreds of transcripts after the onset of zygotic transcription. (a) Plots show RMA-normalized signal intensity of all genes in *pum* mutant versus wild-type embryos at the indicated time intervals. Each dot represents a transcript defined as expressed in at least one time point and in at least one genotype, as described in the Materials and Methods. mRNAs that were at least 1.5 fold up or downregulated with an FDR of < 5% in *pum* mutant embryos are indicated in red or blue, respectively. Dashed lines indicate 1.5-fold increase or decrease in expression, while the solid diagonal line represents no change. N=3 for time intervals 0-1, 1-2, 2-3 and 3-4 hours, while N=2 for the 4-5 h interval. (b) Histogram showing the number of genes significantly upregulated in *pum* mutant embryos, as determined in (a).

In addition to *smg*, the *bard* mRNA and Bard protein are cleared at the end of the MZT (Cao *et al*. 2020; Cao *et al*. 2022). Bard is an F-box protein that acts as a timer for SCF function in clearing SMG protein (Cao *et al*. 2020; Cao *et al*. 2022). Given that the *bard* mRNA is bound by PUM (Laver *et al*. 2015a) and that its 137 nucleotides long 3’ UTR contains three potential PBEs (RNAcompete ranks 10, 53 and 10, Supplemental Table 3), we queried our microarray data to assess whether *bard* mRNA persisted in *pum* mutant embryos and found that it is indeed stabilized at 3-4 and 4-5 hours compared to wild-type embryos (Fig. 6). Thus, both the *smg* mRNA itself and that of *bard*, which encodes the timer SMG clearance, are targets of PUM and are stabilized in *pum* mutants. The potential implications of PUM-mediated decay of *bard* mRNA are outlined in the Discussion.

**Figure 6.**
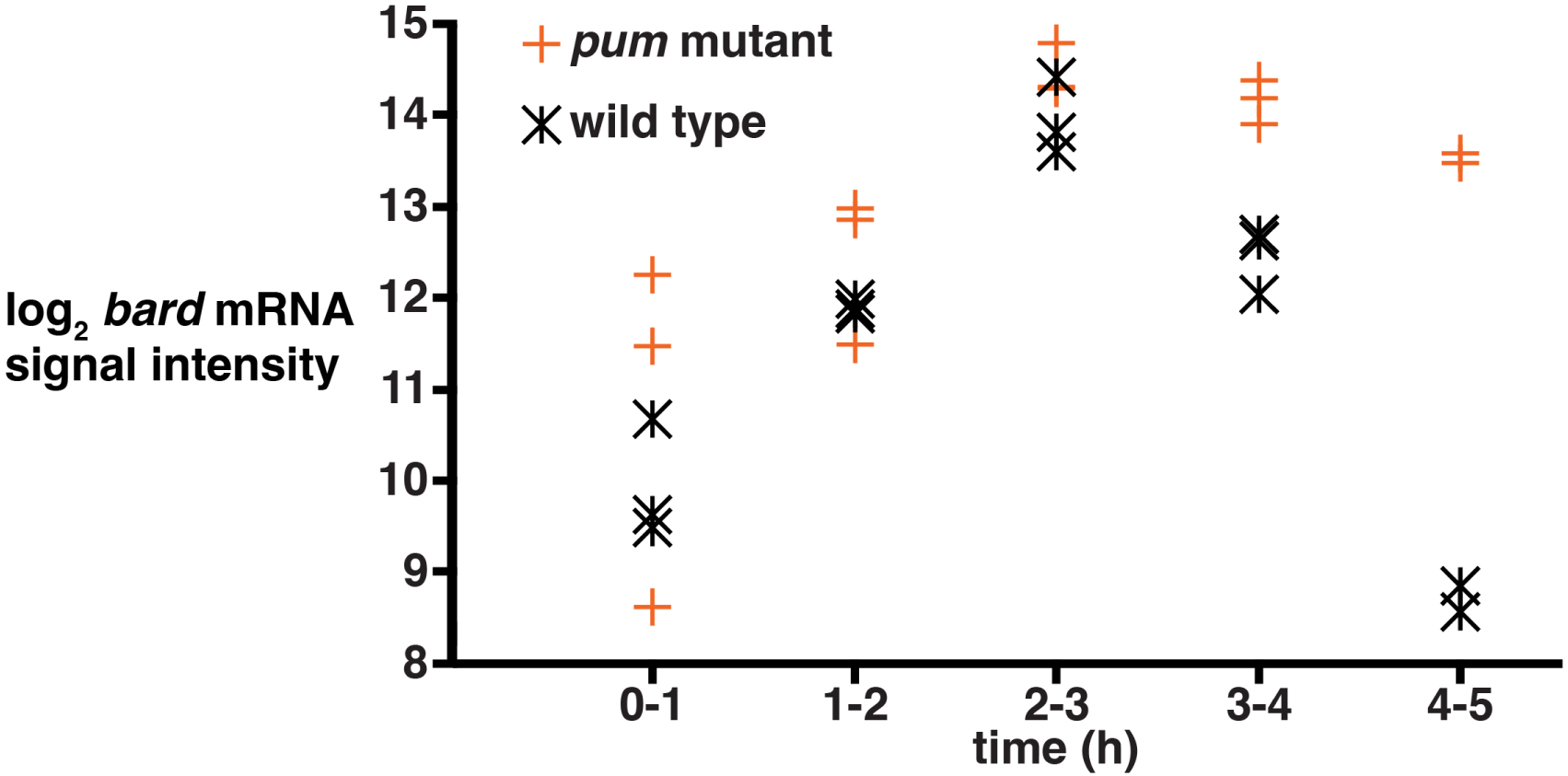
*bard* mRNA is stabilized in *pum* mutant embryos. Plot shows RMA-normalized signal intensity for the *bard* mRNA in *pum* mutant and wild-type embryos at the indicated time intervals. *bard* mRNA in wild-type embryos is expressed in a narrow temporal window, consistent with it being zygotically expressed in the early embryo and degraded at subsequent time points. *bard* mRNA levels are similar in wild-type and *pum* mutant embryos at the first three time points, while being significantly upregulated in *pum* mutant embryos relative to wild type in 3-4 and 4-5 h embryos (FDR < 5%). N=3 for time intervals 0-1, 1-2, 2-3 and 3-4 hours, while N=2 for the 4-5 h interval.

We used two strategies to assess whether the upregulated RNAs were likely to be direct targets of PUM. First, we assessed whether the upregulated RNAs were enriched for predicted PBEs (UGUANA), as determined using RNAcompete (Ray *et al*. 2013) using an approach that incorporates target-site accessibility (i.e., the probability that a PBE is single-stranded; see Materials and methods). The sum of the accessibilities of all PBEs within an RNA was significantly higher for upregulated RNAs compared to co-expressed RNAs that were not upregulated (Wilcoxon Rank Sum test *P*=3.55×10^-6^, Supplemental Fig. 2).

Second, we analyzed three published datasets that identified PUM-bound RNAs in ovaries or early embryos (Gerber *et al*. 2006; Laver *et al*. 2015a; Haugen *et al*. 2024). We found that, in all three cases, RNAs that we identified as upregulated in *pum* mutants were significantly enriched for PUM-bound transcripts (Supplemental Table 4, Fisher’s exact test *P*<10^-6^, odds ratio > 2).

We next asked if the transcripts upregulated in *pum* mutant embryos represent mRNAs that are degraded in wild-type embryos. We made use of data from an earlier analysis that grouped RNAs into five different classes (Thomsen *et al*. 2010): Class I RNAs are stable, Class II RNAs are subject to degradation via maternal factors, Class III RNAs are subject to maternal decay and subsequently accumulate through zygotic transcription, Class IV RNAs are degraded through mechanisms that require zygotic transcription, and Class V RNAs are degraded by both maternal and zygotic pathways. We found that transcripts upregulated in 3-4 and/or 4-5 h *pum* mutant embryos overlapped significantly only with Class IV genes (Fisher’s exact test *P*=5.31×10^-10^, odds ratio=2.05). In contrast, genes from the other four classes were either depleted in or neither enriched nor depleted in *pum* upregulated genes (Supplemental Table 5). These data are consistent with a major role for PUM in an mRNA decay pathway that requires the onset of new transcription in the embryo, as has been previously proposed (De Renzis *et al*. 2007; Benoit *et al*. 2009; Thomsen *et al*. 2010; Laver *et al*. 2015a).

Together our data are consistent with a direct role for PUM in the clearance of a large subset of maternal mRNAs during the late phase of the MZT, a process that requires zygotic transcription.

### BRAT and PUM are likely to function together in late mRNA decay

Our previous genome-wide analysis of BRAT targets showed that *smg* mRNA is both bound by BRAT and stabilized in embryos from *brat* mutant mothers (hereafter referred to as *brat* mutant embryos) relative to wild-type embryos (Laver *et al*. 2015a). We confirmed the latter using RT-qPCR (Fig. 7a). BRAT binds an RNA motif with the core sequence UGUU (Laver *et al*. 2015a; Loedige *et al*. 2015). As described above for PUM and PBEs, we identified UGUU-containing 7mers in the *smg* 3’ UTR and used RNAcompete data to compare their relative affinities for BRAT. One such site had an RNAcompete rank of 4, six had ranks between 16 and 48, and eight had ranks between 64 and 295 (Supplemental Fig. 3 and Supplemental Table 6). Consistent with BRAT playing a role in clearing *smg* mRNA from the embryo, we also detected ectopic SMG protein in 3-4 and 4-5 h *brat* mutant embryos (Fig. 7b and c).

**Figure 7.**
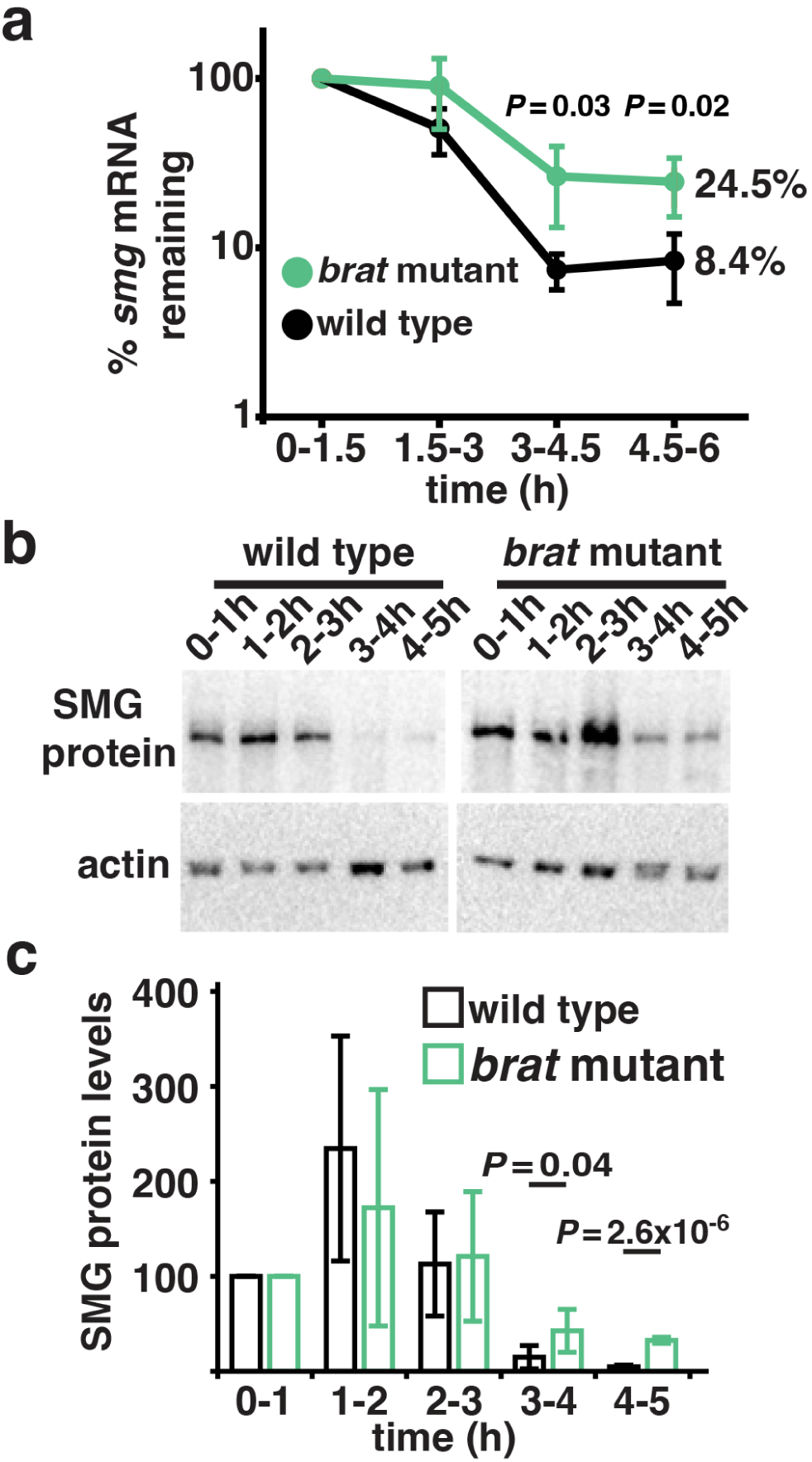
BRAT regulates *smg* mRNA stability and prevents ectopic SMG protein expression. (a) Total RNA was harvested from wild-type and *brat* mutant embryos at the indicated time intervals, and levels of *smg* mRNA were assayed using RT-qPCR. After normalizing *smg* mRNA levels to *RpL32* mRNA, the amount in 0-1.5 h embryos was set to 100%. N=3, with the percent remaining of each mRNA at the last time point indicated. One-tailed Student’s *t*-test *P* values assess the significance of the difference in the levels of *smg* mRNA in wild-type versus *brat* mutant embryos at the last two time points. (b) Total protein was harvested from wild-type and *brat* mutant embryos at the indicated time intervals, and levels of SMG protein were assayed via western blot. Blotting for actin served as the loading control. (c) After normalizing SMG protein levels to the loading control, the amount in 0-1 h embryos was set to 100%. N=4, with error bars represent standard deviation. One-tailed Student’s *t*-test *P* values assess the significance of the difference in the levels of SMG protein in wild-type versus *brat* mutant embryos at the last two time points.

We previously showed that there is a significant overlap between PUM and BRAT bound mRNAs (Laver *et al*. 2015a). We, therefore, expanded our analyses transcriptome-wide to assess the scale of the overlap between BRAT and PUM-mediated mRNA decay. We previously defined three classes of maternally expressed genes (A, B and C) that are upregulated in *brat* mutant embryos and are enriched in BRAT-bound transcripts (Laver *et al*. 2015a). Of these three classes, only C degrades exclusively through a late-acting mechanism, similar to the behaviour of the vast majority of *pum* upregulated genes. Comparison of the class C *brat* genes with the union of genes that are upregulated in 3-4 h and 4-5 h *pum* mutant embryos revealed a significant overlap (Fisher’s exact test *P*=2.16×10^-70^, odds ratio=15.01, Supplemental Table 7). Taken together, these data are consistent with PUM and BRAT functioning together to degrade transcripts during the MZT.

### Ectopic expression of SMG protein disrupts the transcriptome of *pum* and *brat* mutant embryos

Given SMG’s major role in maternal transcript clearance during the MZT (Tadros *et al*. 2007) and the evidence that persistent SMG downregulates several zygotically expressed transcripts containing SREs (Cao *et al*. 2020), we reasoned that ectopic SMG in *pum* and *brat* mutant embryos would globally downregulate SRE-containing mRNAs. To assess this, we identified genes whose expression decreases significantly late in the MZT in *pum* or *brat* mutants compared to wild type.

In 3-4 and 4-5 h *pum* mutant embryos we detected 506 and 1517 downregulated genes respectively (Fig. 5a and Fig. 8a, Supplemental File 1; FDR of < 5% and a fold change of < −1.5). These genes significantly overlapped (Fisher’s exact test 4.11×10^-139^, odds ratio=12.55, Supplemental Table 8) with the union of these two lists, totalling 1625 genes. In contrast, during the first three hours of embryogenesis only 14 genes were downregulated in *pum* mutant embryos (one of which was the *pum* transcript itself, as expected since the *pum^Msc^* allele likely fails to produce *pum* mRNA).

**Figure 8.**
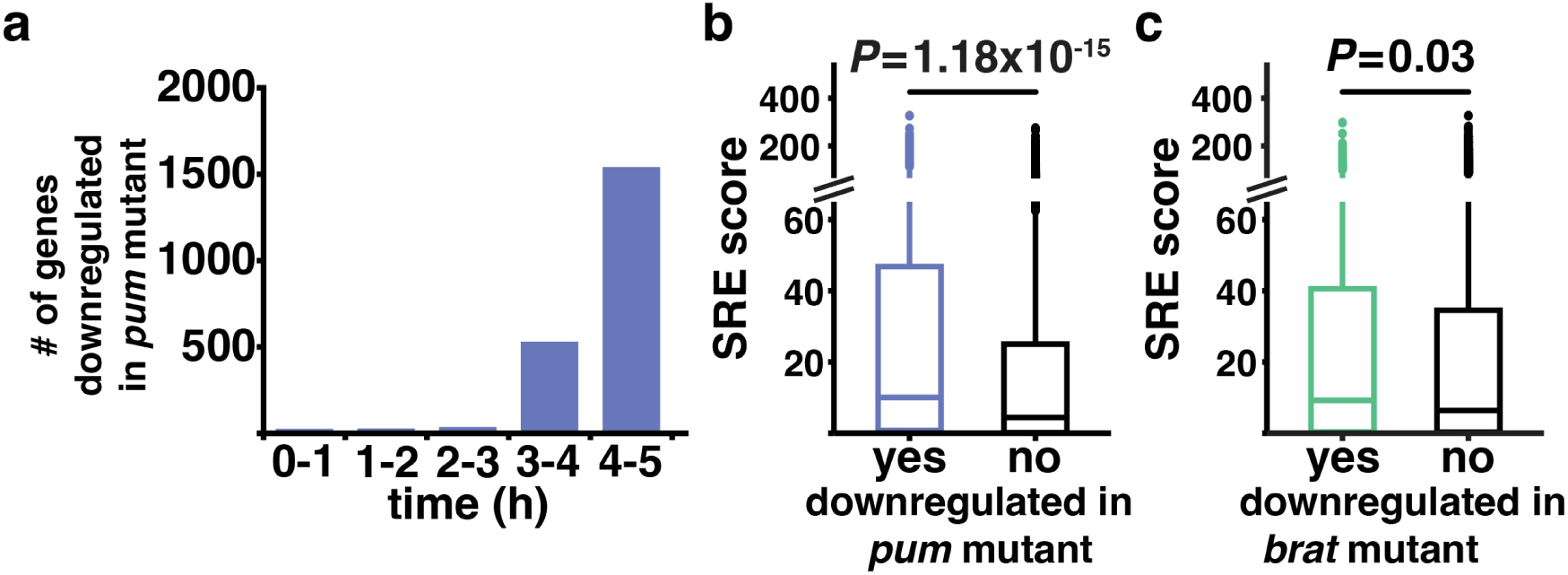
Ectopic expression of SMG protein disrupts the transcriptome of *pum* and *brat* mutant embryos. (a) Histogram showing the number of genes significantly downregulated in *pum* mutant embryos, as determined in Fig. 5a. Box and whisker plots comparing the SRE scores of downregulated transcripts versus all other expressed transcripts in 3-4 and/or 4-5 h *pum* mutant embryos (b) or 3-6 h *brat* mutant embryos (c). The Wilcoxon Rank Sum test *P* values are indicated.

According to our hypothesis, the downregulated transcripts should be enriched for The SRE score for an RNA is the sum of the probabilities that each CNGGN_0-4_ within a transcript will form an SRE stem/loop (Chen *et al*. 2014; Siddiqui *et al*. 2024). We found that the SRE scores for the RNAs downregulated in 3-4 and 4-5 h *pum* mutant embryos were significantly higher than for RNAs that were not downregulated (Fig. 8b, Wilcoxon Rank Sum test *P*=1.18×10^-15^).

Our previous transcriptomic analysis of *brat* mutant embryos identified 527 downregulated genes in 3-6 h *brat* mutant embryos (Laver *et al*. 2015a). We found that these genes downregulated genes are enriched in SREs (Fig 8c, Wilcoxon Rank Sum test *P*=0.03). Furthermore, these genes significantly overlapped with those downregulated in 3-4 and 4-5 h *pum* mutant embryos (Fisher’s exact *P*=1.38×10^-52^, odds ratio=4.93, Supplemental Table 9). This is consistent with the possibility that the same mechanism – targeting by SMG – underlies the downregulation of genes in both *brat* and *pum* mutant embryos.

In summary, our data support a model whereby ectopic SMG protein in *pum* and *brat* mutant embryos targets hundreds of transcripts for degradation, thereby substantially disrupting the embryonic transcriptome.

## DISCUSSION

mRNA decay during the MZT occurs in temporal waves, which can be divided into those that occur early in embryogenesis and rely solely on maternally provided products, while later phases require zygotically expressed factors (Vastenhouw *et al*. 2019; Harrison *et al*. 2023). Key to understanding these waves is identifying the *trans*-acting factors involved. Here we have shown that PUM functions in an mRNA decay pathway that requires zygotic transcription and that many of the affected mRNAs are also targeted by BRAT. PUM’s function contrasts with that of BRAT, as the latter also functions in a PUM-independent decay pathway that acts earlier and does not require new transcription.

We have found that the *smg* mRNA is a direct target of PUM- and, likely, BRAT-mediated degradation. Ectopic expression of SMG protein in *pum* or *brat* mutants leads to downregulation of SRE-containing transcripts, thereby disrupting the embryonic transcriptome. In wild-type embryos, SMG protein is highly expressed in a narrow window over the first three hours of embryogenesis. First, *smg* mRNA is repressed during oogenesis and derepressed in early embryos (Tadros *et al*. 2007). Second, the SCF E3 ligase clears SMG protein at the end of the MZT (Cao *et al*. 2020). Targeting of SMG by SCF requires zygotic transcription and translation of the *bard* mRNA, which encodes an F-box protein that binds SMG and triggers its proteolysis (Cao *et al*. 2022). We have previously shown that persistent SMG downregulates several SRE-containing transcripts (Cao *et al*. 2020). Here we have shown that ectopic SMG leads to global downregulation of a subset of the transcriptome that is enriched for SREs. This confirms that SMG clearance is essential for an orderly MZT as previously hypothesized (Cao *et al*. 2020).

In those earlier experiments, while *smg* mRNA clearance was normal, the SMG protein was stabilized by deletion of the domain that interacts with the SCF E3 ligase. Here we have shown that stabilization of the *smg* RNA also leads to persistence of SMG protein despite its interaction with SCF. We reconcile these observations as follows: During normal development both the *smg* mRNA and the SMG protein are targeted for clearance – the former by PUM and BRAT and the latter by SCF. If SCF function is abrogated in the context of normal clearance of *smg* mRNA (Cao *et al*. 2020) then SMG protein persists. However, if, as in the current study, SCF function is retained but *smg* mRNA persists, then ongoing translation of new SMG protein results in ectopic SMG.

A further possible complexity is introduced by our analyses of the *bard* mRNA, which encodes an F-box protein that acts as a timer for SCF function in clearing SMG (Cao *et al*. 2020; Cao *et al*. 2022), and is bound by PUM (Laver *et al*. 2015a). We have shown here that *bard* mRNA has potential PBEs in its 3’ UTR and its levels are significantly increased in 3-4-and 4-5 h *pum* mutant embryos. In contrast to the *smg* mRNA, the *bard* mRNA is neither bound by BRAT nor upregulated in *brat* mutant embryos (at the time it was listed as CG14317) (Laver *et al*. 2015a), and it also lacks BRAT-binding sites in its 3’ UTR (i.e., no UGUU motifs). Thus, clearance of *bard* mRNA is PUM-dependent but BRAT-independent. In the absence of PUM, the resulting ectopic Bard may limit the levels of ectopic SMG seen in mutants since SCF targeting of SMG protein would be predicted to persist (Cao *et al*. 2020). In summary, during normal development PUM functions both to clear the *smg* mRNA and, consequently, prevent synthesis of new SMG protein. PUM also promotes clearance of *bard* mRNA which encodes the timer of SCF-dependent elimination of SMG and future analyses will focus on the functional significance of Bard clearance.

We do not know why PUM’s role in maternal transcript clearance is restricted to the late MZT, despite the protein being present at similar levels throughout the MZT (Macdonald 1992; Casas-Vila *et al*. 2017; Cao *et al*. 2020). Our data argue against a model that proposes temporally regulated PUM recruitment to its targets only after the onset of zygotic transcription since we see similar levels of *smg* mRNA enrichment in PUM IPs on 0-2 h and 2-4 h embryos (Fig. 3a). At this time, we do not know whether this will generalize to many or even all PUM’s target transcripts.

Here we discuss two hypotheses to explain the timing of *smg* mRNA degradation by PUM. First, it is possible that zygotically expressed factors post-translationally modify PUM thus altering its regulatory role. For example, PUM is known to function through the recruitment of the CCR4/NOT deadenylase complex, resulting in the removal of the mRNA’s poly(A) tail, which in turn can trigger transcript decay (Weidmann *et al*. 2014; Arvola *et al*. 2020; Haugen *et al*. 2022; Haugen *et al*. 2024). Post-translational modifications could alter PUM’s ability to recruit these factors. This type of mechanism and the one discussed below, could also apply to BRAT, but would have to account for the fact that the degradation of some BRAT targets requires new transcription, while clearance of other BRAT targets does not.

Second, and not mutually exclusive, zygotically synthesized factors may act together with PUM to trigger target decay during the late phase of the MZT. Previous work has shown that PUM can use distinct cofactors to regulate different targets (Arvola *et al*. 2017; Malik *et al*. 2019). For example, posterior regulation of *hb* requires both BRAT and Nanos, whereas posterior regulation of *cyclin B* requires PUM and Nanos but not BRAT (Murata and Wharton 1995; Asaoka-Taguchi *et al*. 1999; Sonoda and Wharton 1999; Sonoda and Wharton 2001; Kadyrova *et al*. 2007). Moreover, because Nanos is absent outside the posterior, PUM-mediated regulation of *cyclin B* in the bulk of the embryo is likely Nanos-independent (Vardy and Orr-Weaver 2007). Similarly, PUM-dependent regulation of *paralytic* mRNA requires Nanos and BRAT in motor neurons, but in most of the CNS it requires Nanos without BRAT (Muraro *et al*. 2008).

Finally, the work presented here supports a model where SMG is involved in a negative autoregulatory feedback loop that temporally restricts its own expression during the MZT (Fig. 9). This is accomplished in two ways: First, Bard, which is zygotically expressed, triggers SMG protein degradation. Second, an as-yet-unidentified zygotic factor(s) functions together with PUM and BRAT to trigger *smg* mRNA decay. Thus, clearance of both *smg* mRNA and SMG protein requires zygotic transcription. Given that SMG is required for robust zygotic RNA synthesis (Benoit *et al*. 2009; Luo *et al*. 2016), SMG is indirectly involved in its own clearance during the MZT.

**Figure 9.**
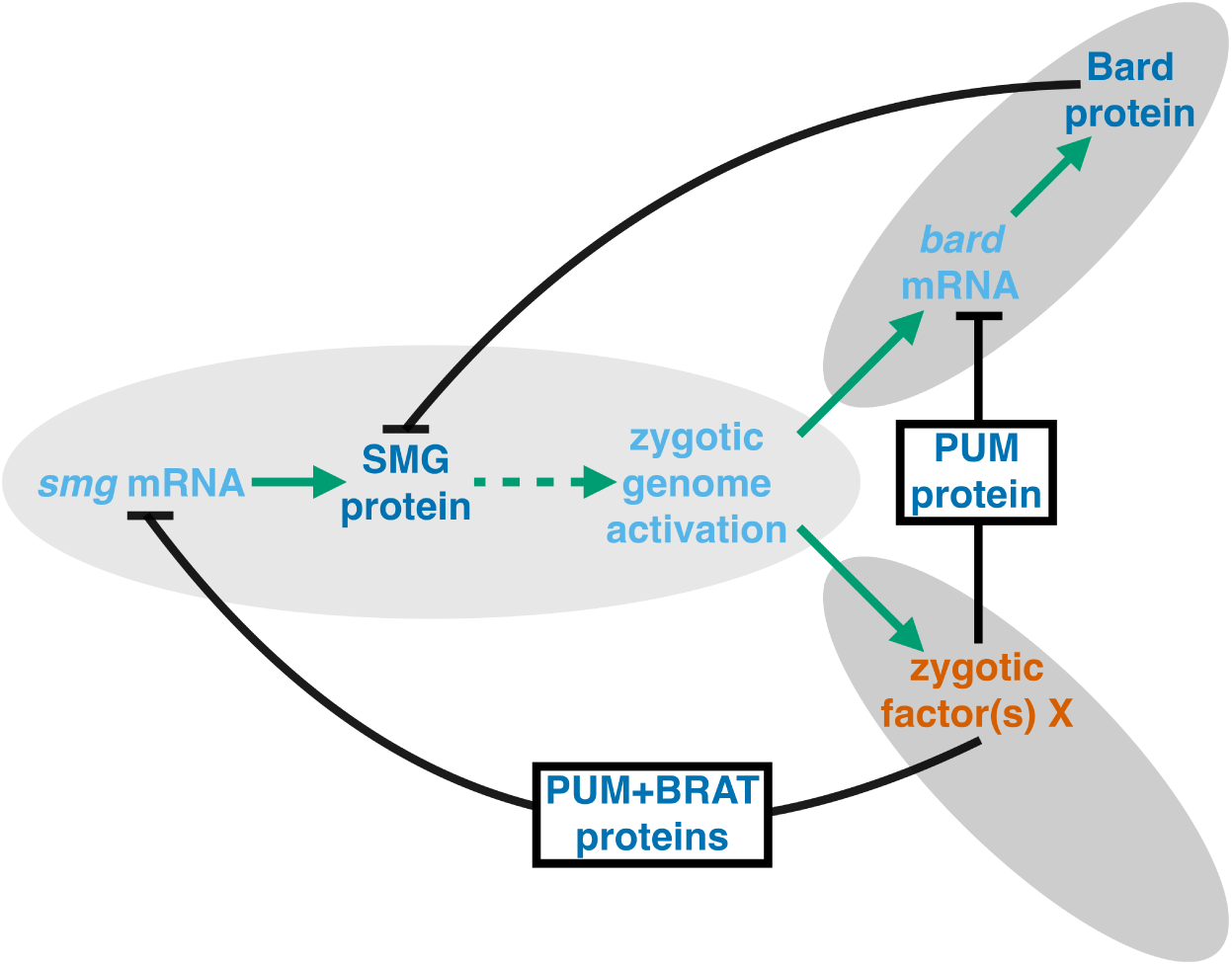
A proposed model for an autoregulatory feedback loop that clears SMG protein during the MZT. The translational activation of *smg* mRNA in newly-laid embryos results in the accumulation of SMG protein (Tadros *et al*. 2007). SMG protein, through an unknown mechanism that presumably relies on SMG-mediated regulation of one or more target transcripts, is required for high-level zygotic transcription (Benoit *et al*. 2009; Luo *et al*. 2016). This new transcription includes the *bard* mRNA and the resulting Bard protein triggers the degradation of the SMG protein (Cao *et al*. 2020; Cao *et al*. 2022). New transcription also results in the production of an as-yet-unidentified factor(s) (X) that collaborates with PUM and BRAT to trigger *smg* mRNA degradation. PUM-mediated degradation of *bard* mRNA may also be required to temporally regulate the expression of Bard protein.

## DATA AVAILIBILTY STATEMENT

Reagents generated in this study are available from the authors upon reasonable request. The classes of mRNAs defined in Thomsen et al. (2010) can be found at ArrayExpress (E-MEXP-2580). The remaining data sets are available at GEO. GSE48645 and GSE65661 characterize the transcriptomes of *pum* mutant data (this study) and *brat* mutant embryos (Laver *et al*. 2015a), respectively. GSE3582 (Gerber *et al*. 2006), GSE60466 (Laver *et al*. 2015a), and GSE240494 (Haugen *et al*. 2024), identify PUM bound mRNAs. GSE41235 (Ray *et al*. 2013) and GSE60498 (Laver *et al*. 2015a) include the RNAcompete data for PUM and BRAT, respectively. The code for MFRA can be found at https://github.com/LipshitzLabToronto. The authors affirm that all of the other data necessary for confirming the conclusions of the article are present within the article, figures, tables, and supplemental files.

## SUPPLEMENTAL FIGURES AND TABLES

**Supplemental Figure 1.**
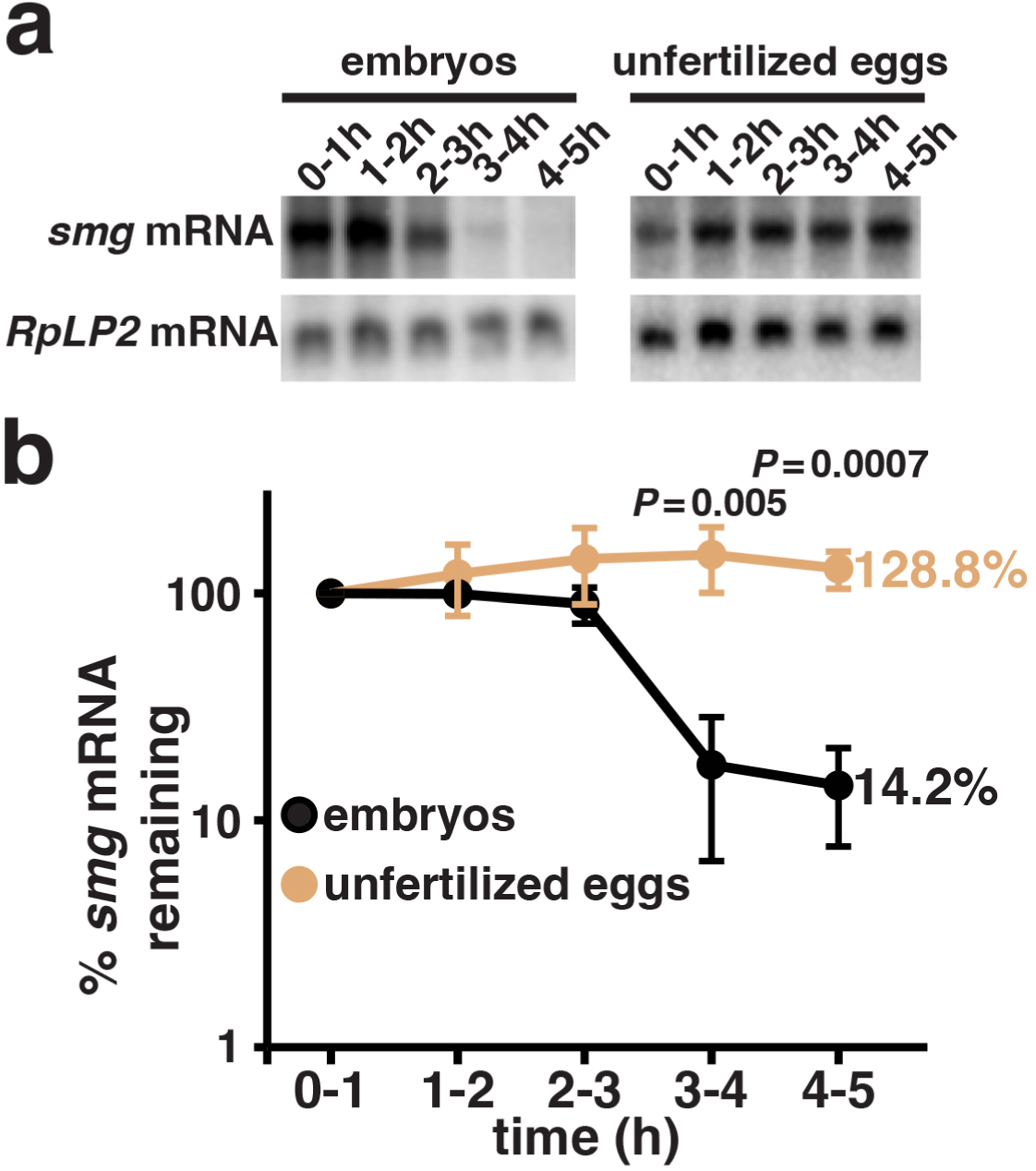
s*m*g mRNA is stable in unfertilized eggs. (a) Total RNA was harvested from wild-type embryos or unfertilized eggs at the indicated time intervals and subjected to northern blot analysis using *smg* or *RpLP2* mRNA probes. (b) After normalizing *smg* mRNA levels to *RpLP2* levels, the amount in the first time point was set to 100%. N=3, error bars represent standard deviation, while the percent remaining of each mRNA at the last time point is indicated. One-tailed Student’s *t*-test *P* values assessing the significance of the difference in the levels of *smg* mRNA in embryos versus unfertilized eggs at the last two time points are indicated.

**Supplemental Figure 2.**
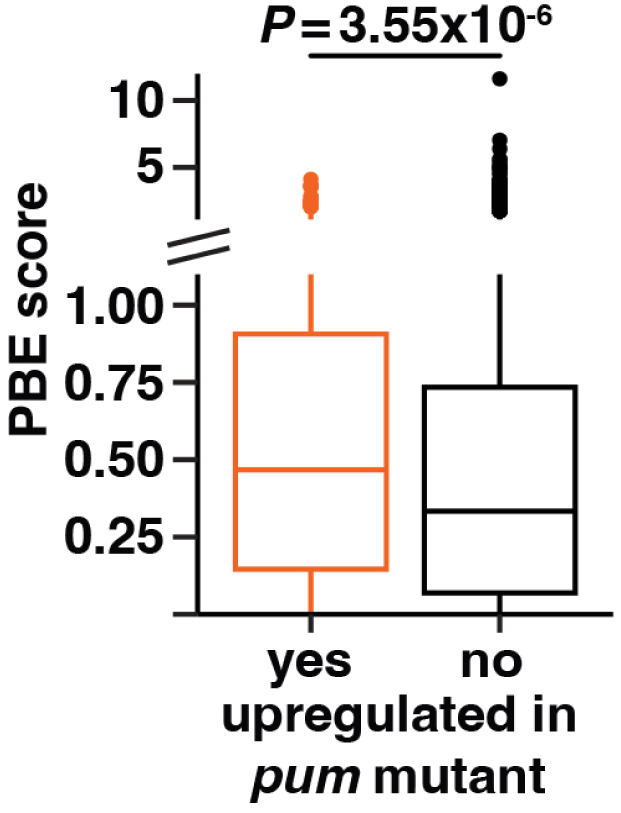
Genes upregulated in *pum* mutant embryos are enriched in predicted PBEs. A box and whisker plot comparing the PBE score, which is the sum of the accessibilities of all UGUANA sequences within an mRNA, of transcripts upregulated in 3-4 and/or 4-5 h *pum* mutant embryos versus all other expressed transcripts. The Wilcoxon Rank Sum test *P* value is indicated.

**Supplemental Figure 3.**
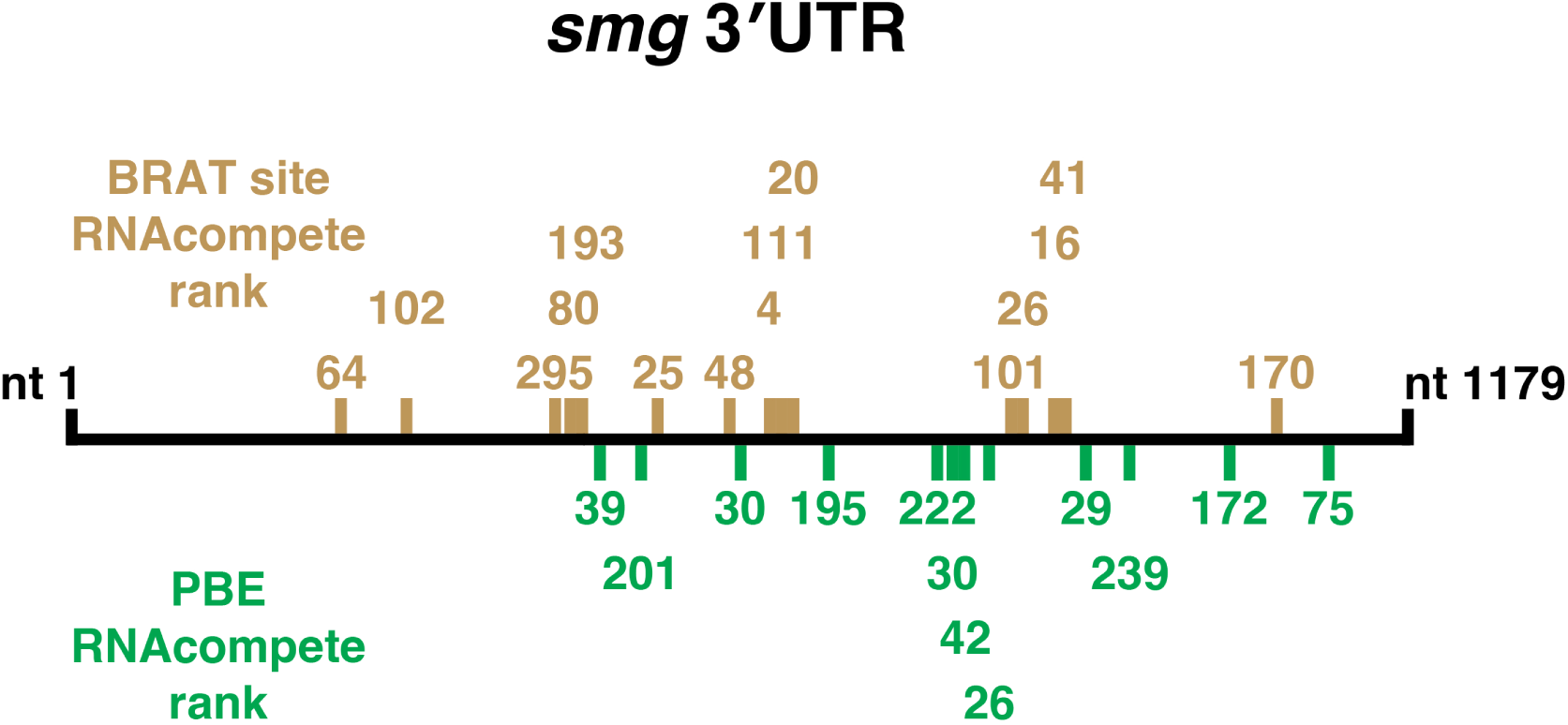
The location of potential PBEs and BRAT binding sites in the *smg* 3’ UTR. The *smg* 3’ UTR is diagrammed, indicating the location of potential PBEs, using the sequence UGUANNNN and BRAT binding sites containing the core BRAT motif UGUU. The rank of each site in the RNAcompete data is indicated, with rank 1 corresponding to the site with the highest apparent affinity for the indicated protein.

**Supplemental Table 1.**
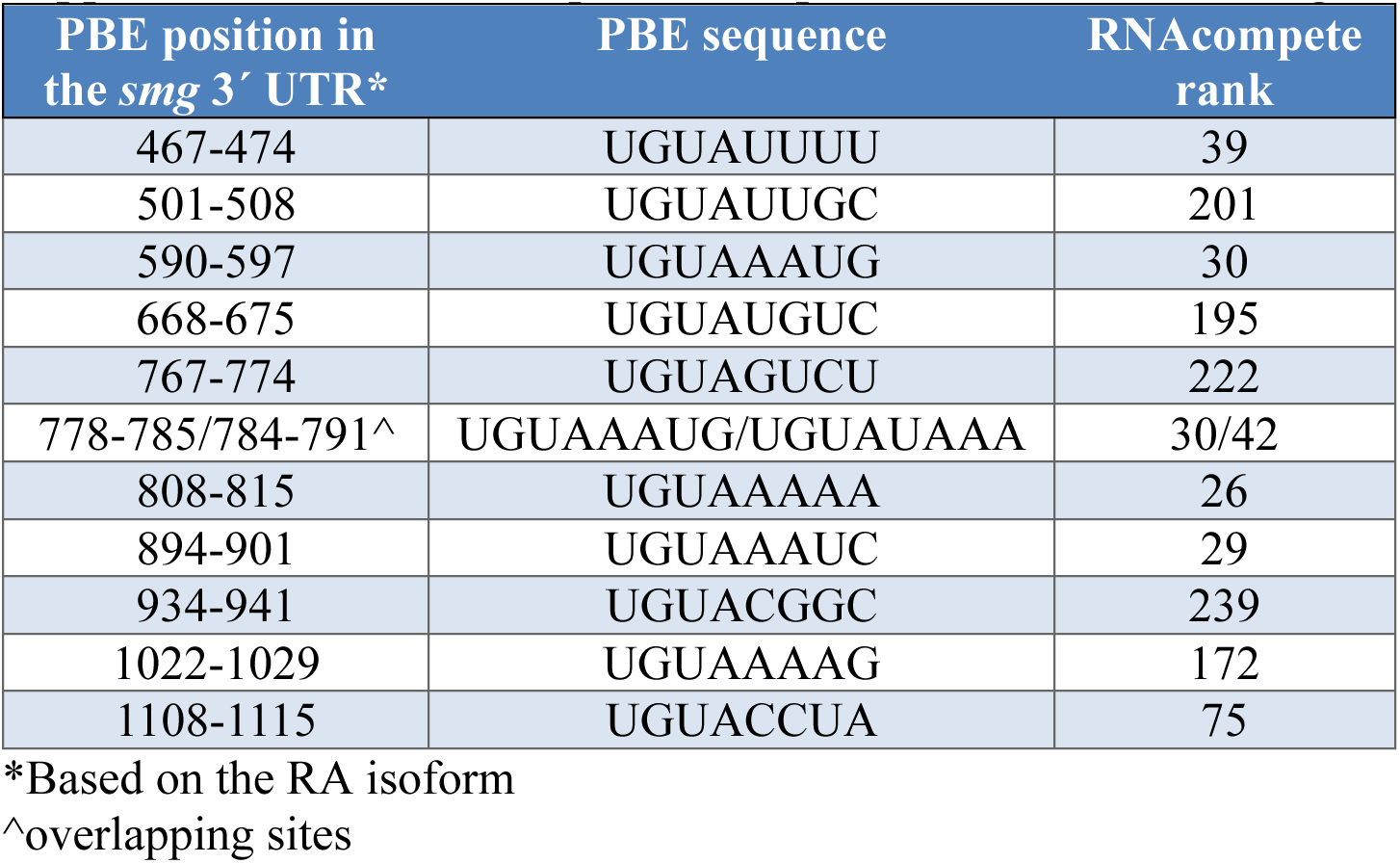
The position of predicted PBEs in the *smg* mRNA’s 3’ UTR.

| PBE position in the *smg* 3' UTR* | PBE sequence | RNAcomplete rank |
| --- | --- | --- |
| 467-474 | UGUAUUUU | 39 |
| 501-508 | UGUAUUGC | 201 |
| 590-597 | UGUAAAUG | 30 |
| 668-675 | UGUAUGUC | 195 |
| 767-774 | UGUAGUCU | 222 |
| 778-785/784-791^ | UGUAAAUG/UGUAUAAA | 30/42 |
| 808-815 | UGUAAAAA | 26 |
| 894-901 | UGUAAAUC | 29 |
| 934-941 | UGUACGGC | 239 |
| 1022-1029 | UGUAAAAG | 172 |
| 1108-1115 | UGUACCUA | 75 |

\*Based on the RA isoform
^overlapping sites

**Supplemental Table 2.**
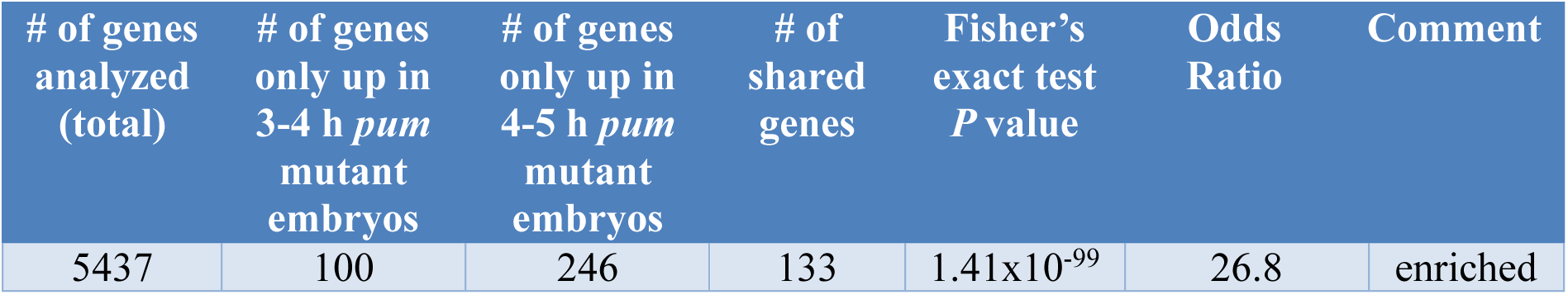
Overlap of upregulated genes in 3-4 and 4-5 h *pum* mutant embryos.

| # of genes analyzed (total) | # of genes only up in 3-4 h *pum* mutant embryos | # of genes only up in 4-5 h *pum* mutant embryos | # of shared genes | Fisher's exact test *P* value | Odds Ratio | Comment |
| --- | --- | --- | --- | --- | --- | --- |
| 5437 | 100 | 246 | 133 | 1.41x10<sup>-99</sup> | 26.8 | enriched |

**Supplemental Table 3.** The position of potential PBEs in the *bard* mRNA’s 3’ UTR.

| PBE motif position in the *bard* 3' UTR* | motif sequence | RNAcomplete rank |
| --- | --- | --- |
| 26-33 | UGUAUAUA | 10 |
| 65-72 | UGUAAUUU | 53 |
| 87-94 | UGUAAUUU | 10 |

\*Based on the RA isoform

**Supplemental Table 4.**
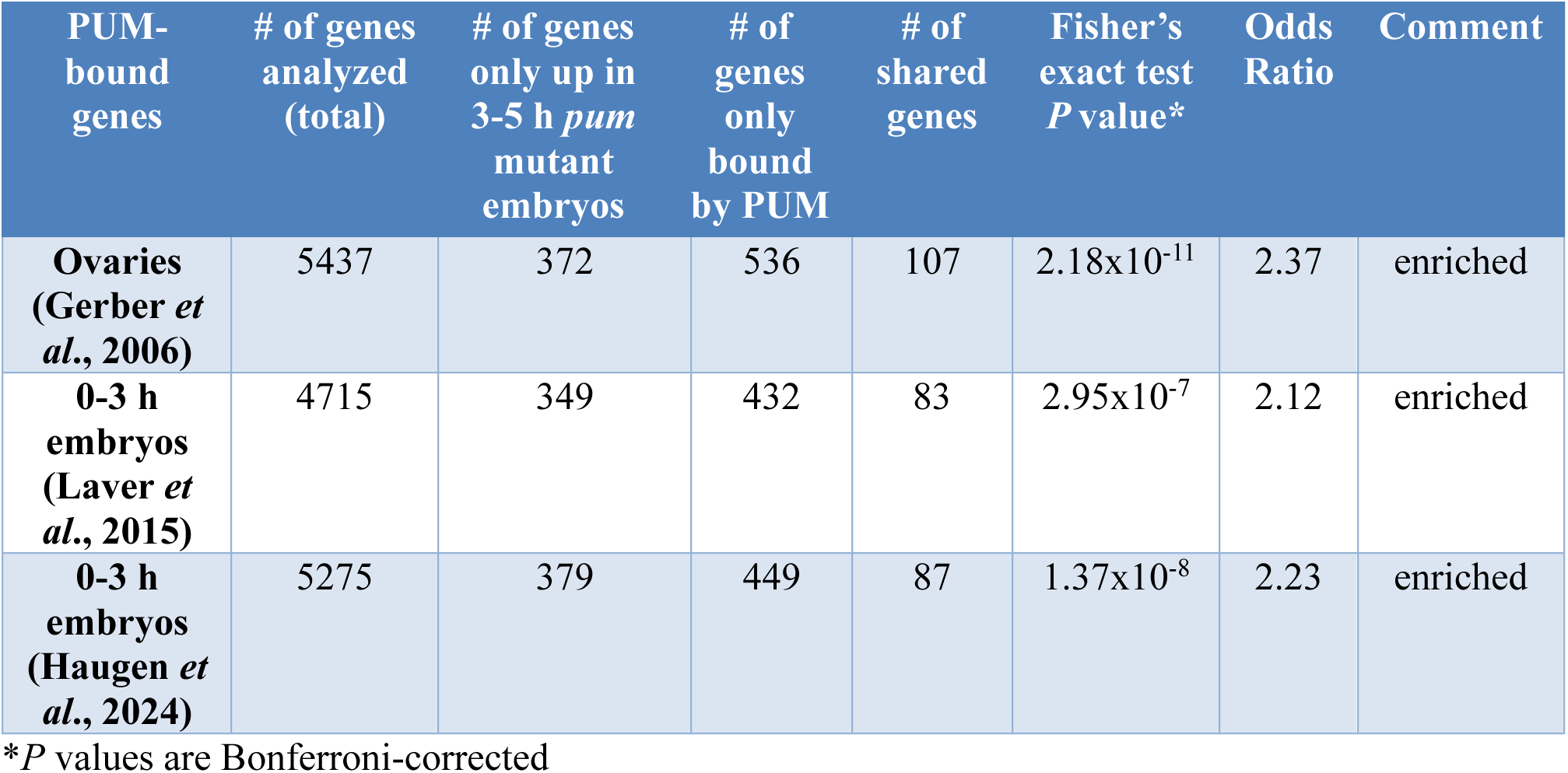
Overlap of genes upregulated in a *pum* mutant relative to PUM-bound genes.

| PUM-bound genes | # of genes analyzed (total) | # of genes only up in 3-5 h *pum* mutant embryos | # of genes only bound by PUM | # of shared genes | Fisher's exact test *P* value* | Odds Ratio | Comment |
| --- | --- | --- | --- | --- | --- | --- | --- |
| Ovaries (Gerber *et al.*, 2006) | 5437 | 372 | 536 | 107 | 2.18x10<sup>-11</sup> | 2.37 | enriched |
| 0-3 h embryos (Laver *et al.*, 2015) | 4715 | 349 | 432 | 83 | 2.95x10<sup>-7</sup> | 2.12 | enriched |
| 0-3 h embryos (Haugen *et al.*, 2024) | 5275 | 379 | 449 | 87 | 1.37x10<sup>-8</sup> | 2.23 | enriched |

\**P* values are Bonferroni-corrected

**Supplemental Table 5.** Overlap of genes upregulated in a *pum* mutant relative to the classes of genes defined in Thomsen *et al*., (2010).

| Thomsen et al., Class | # of genes analyzed (total) | # of genes only in the Thomsen class | # of genes only up-regulated in *pum* mutant | # of shared genes | Fisher's exact test *P* value* | Odds Ratio | Comment |
| --- | --- | --- | --- | --- | --- | --- | --- |
| I | 4019 | 1052 | 279 | 126 | 1 | 1.1 | Neither enriched nor depleted |
| II | 4019 | 281 | 396 | 9 | 3.09x10<sup>-5</sup> | 0.27 | Depleted |
| III | 4019 | 284 | 402 | 3 | 1.50x10<sup>-9</sup> | 0.09 | Depleted |
| IV | 4019 | 907 | 240 | 165 | 5.31x10<sup>-10</sup> | 2.05 | Enriched |
| V | 4019 | 1146 | 294 | 111 | 0.40 | 0.81 | Neither enriched nor depleted |

\**P* values are Bonferroni-corrected

**Supplemental Table 6.** The position of potential BRAT-binding sites in the *smg* mRNA’s 3’ UTR.

| BRAT motif position in the *smg* 3' UTR* | motif sequence | RNAcomplete rank |
| --- | --- | --- |
| 238-244 | UUGUUUG | 64 |
| 296-302 | UGUUUAU | 102 |
| 426-432 | CAAUGUU | 295 |
| 437-443/440-446^ | UGUUUGU/ UUGUGUU | 80/193 |
| 561-567 | UUAUGUU | 25 |
| 580-586 | UUGUUUA | 48 |
| 617-623/622-628/628-634^ | UGUUAUG/ UGUUUUA/ AGUUGUU | 4/111/20 |
| 829-835/832-838^ | UGUUUUU/ UUUUGUU | 101/26 |
| 866-872/870-876^ | AUUUGUU/ GUUUGUU | 16/41 |
| 1066-1072 | AAAUGUU | 170 |

\*Based on the RA isoform
^overlapping sites

**Supplemental Table 7.** Overlap of genes upregulated in *pum* mutant embryos versus *brat* mutant class C genes (Laver *et al*., 2015).

| # of genes analyzed (total) | # of genes up in only 3-5 h *pum* mutant embryos | # of genes only in BRAT class C | # of shared genes | Fisher's exact test *P* value | Odds Ratio | Comment |
| --- | --- | --- | --- | --- | --- | --- |
| 3944 | 246 | 117 | 125 | 2.16x10<sup>-70</sup> | 15.0 | enriched |

**Supplemental Table 8.**
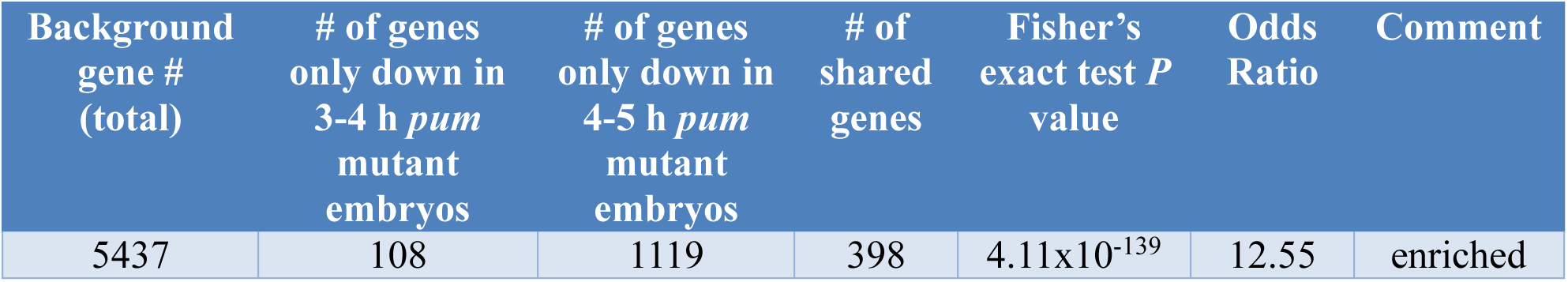
Overlap of downregulated genes in 3-4 and 4-5 h *pum* mutant embryos.

**Supplemental Table 9.** Overlap of genes downregulated in *pum* mutant embryos versus *brat* mutant embryos (Laver *et al*., 2015).

| Background gene # (total) | # of genes only down in 3-5 h *pum* mutant embryos | # of genes only down in 3-6 h *brat* mutant embryos | # of shared genes | Fisher's exact test *P* value | Odds Ratio | Comment |
| --- | --- | --- | --- | --- | --- | --- |
| 3944 | 848 | 170 | 267 | 1.38x10<sup>-52</sup> | 4.93 | enriched |

